# A versatile framework for drug-target interaction prediction by considering domain specific features

**DOI:** 10.1101/2023.08.01.551396

**Authors:** Shuo Liu, Jialiang Yu, Ningxi Ni, Zidong Wang, Mengyun Chen, Yuquan Li, Chen Xu, Yahao Ding, Jun Zhang, Xiaojun Yao, Huanxiang Liu

## Abstract

Predicting drug-target interaction (DTI) is a critical and rate-limiting step in drug discovery. Traditional wet-lab experiments are reliable but expensive and time-consuming. Recently, deep learning has revealed itself as a new and promising tool for accelerating the DTI prediction process because its powerful performance. Due to the vast chemical space, the DTI prediction models are typically expected to discover drugs or targets that are absent from the training set. However, generalizing prediction performance to novel drug-target pairs that belong to different distributions is a challenge for deep learning methods. In this work, we propose an Ensemble of models that capture both Domain-generIc and domain-Specific features (E-DIS) to learn diversity domain features and adapt to out-of-distribution (OOD) data. We employed Mixture-of-Experts (MOE) as a domain-specific feature extractor for the raw data to prevent the loss of any crucial features by the encoder during the learning process. Multiple experts are trained on different domains to capture and align domain-specific information from various distributions without accessing any data from unseen domains. We evaluate our approach using four benchmark datasets under both in-domain and cross-domain settings and compare it with advanced approaches for solving OOD generalization problems. The results demonstrate that E-DIS effectively improves the robustness and generalizability of DTI prediction models by incorporating diversity domain features.

## 1 Introduction

The accurate prediction of DTI is a basic task in the field of new drug design and development, with significant implications for drug repositioning, drug discovery, side-effect prediction and drug resistance [1–3]. While traditional wet-lab experiments are reliable, they are also costly and time-consuming [4]. Deep learning has recently been widely used to accelerate the DTI prediction process due to its fast and powerful learning ability, which allows compounds with high affinity to be quickly selected from large amounts of data for further study [5]. The drug-target pairs that need to be predicted in real-world applications are often unseen in the training data due to the vast chemical space [6]. However, the performance of deep learning methods decreases rapidly when predicting novel drug-target pairs. Generalizing prediction performance to OOD data is a challenging for deep learning methods [7,8].

To overcome these challenge, researchers have been working on developing models with strong generalization ability to accurately predict drug-target pairs that are unseen in the training data [9,10]. One of the most notable approaches is to learn domain-generic features, with the assumption that these generic representations exist across all data instances [11–13]. Adversarial domain adaptation is a method that aims to learn domain-generic features by minimizing the distribution discrepancy across domains [14]. However, deep learning methods can sometimes rely too heavily on a single feature, leading to shortcuts that hinder accurate predictions on OOD data [15]. To address this issue, Pagliardini et al. proposed a technique that captures a diverse set of domain-generic features by enforcing agreement among models on the source domain while introducing disagreement on the target domain [16]. Nonetheless, some studies[13,17,18] have shown that forcing the model to learn domain-generic features on domains with vastly different marginal label distributions can lead to a deterioration in performance. Hence, learning domain-generic features alone may not always be sufficient for achieving successful generalization.

On the other hand, incorporating domain-specific features can enhance the model’s ability to capture unique patterns and relationships specific to each domain, leading to more accurate predictions on domain-specific data [19]. One popular deep learning technique that leverages this idea is the MOE approach, which has been successfully applied to various tasks [11,20,21]. Each input is assigned to an expert that specializes in handling data from a specific domain. This allows the model to effectively capture domain-specific features and make more precise predictions. Recent studies have demonstrated that MOE exhibits superior generalization ability compared to the original network, especially when the number of source domains increases [7,22–24]. However, no existing research has explored the connection between MOE and DTI prediction.

Here, in order to enhance the generalization ability of DTI prediction model, we proposed a simple yet efficient prediction framework by training Ensembles of Domain-generIc feature extractor and domain-Specific feature extractor (E-DIS). Our approach involves training an ensemble of models that capture both domain-generic and domain-specific features. Initially, we trained a domain-generic feature extractor using all the available training data, and then modeled the potential space for drug-target pairs as a mixture of different domains using a gating network that relied on the domain-generic feature. Unlike other approaches[11,20,21] that used MOE as a layer in the model, we employed MOE as a domain-specific feature extractor for the raw data. This prevents the loss of any crucial features by the encoder during the learning process. E-DIS represents the first attempt to simultaneously learn both domain-generic and domain-specific features for the DTI prediction task. By incorporating diversity domain features, E-DIS avoids relying solely on one feature and mitigates the risk of spurious correlations.

## 2 Methods

### 2.1 Datasets

Four datasets were used to train and validate our model. Table 1 shows the statistics information of the four datasets. The benchmark datasets used for the regression task in this study are the same as those used in GraphDTA[25], including the Davis[26] and KIBA[27] datasets. The binding affinity of the Davis dataset is measured using the dissociation constant (Kd) values, which are transformed into pKd for binding affinity prediction by Equation (1), with values ranging from 5.0 to 10.8. The binding affinity of the KIBA dataset is measured using the KIBA score[27] computed from the combination of heterogenous information sources such as Ki, Kd, and IC50, with values ranging from 0.0 to 17.2. For the classification task, two widely used DTI datasets (BindingDB[28], BIOSNAP[6,29]) are used. BindingDB is a low-bias version of the dataset constructed from BindingDB database by Bai et al,[28] while BIOSNAP is a balanced dataset with the same number of positive and negative samples created from the DrugBank database by Huang et al.[29]

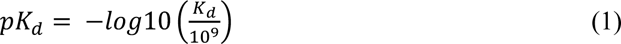

**Table 1.** The statistics of experimental datasets.

| Datasets | Task type | Compounds | Proteins | Interactions |
| --- | --- | --- | --- | --- |
| Davis | Regression | 68 | 442 | 30056 |
| KIBA | Regression | 2111 | 229 | 118254 |
| BindingDB | Classification | 14,643 | 2,623 | 49199 |
| BIOSNAP | Classification | 4,510 | 2181 | 27,464 |

### 2.2 Method overview

Figure 1 illustrates an overview of the E-DIS framework, which involves ensembles of a domain-generic feature extractor and a domain-specific feature extractor. The process began by training the domain-generic feature extractor using the entire training dataset. Subsequently, we employed MOE model, which consisted of a set of expert networks and a trainable gating network, to learn domain-specific features. The gating network routed the input data to the corresponding experts that specialize in specific domain, based on the features learned by the domain-generic feature extractor. Each expert network within the MOE can be any DTI prediction network designed to capture domain-specific features. Finally, we predicted the interaction between a given drug-target pair by computing the weighted average of the predicted values obtained from both the domain-generic feature extractor and the domain-specific feature extractor.

**Figure 1.**
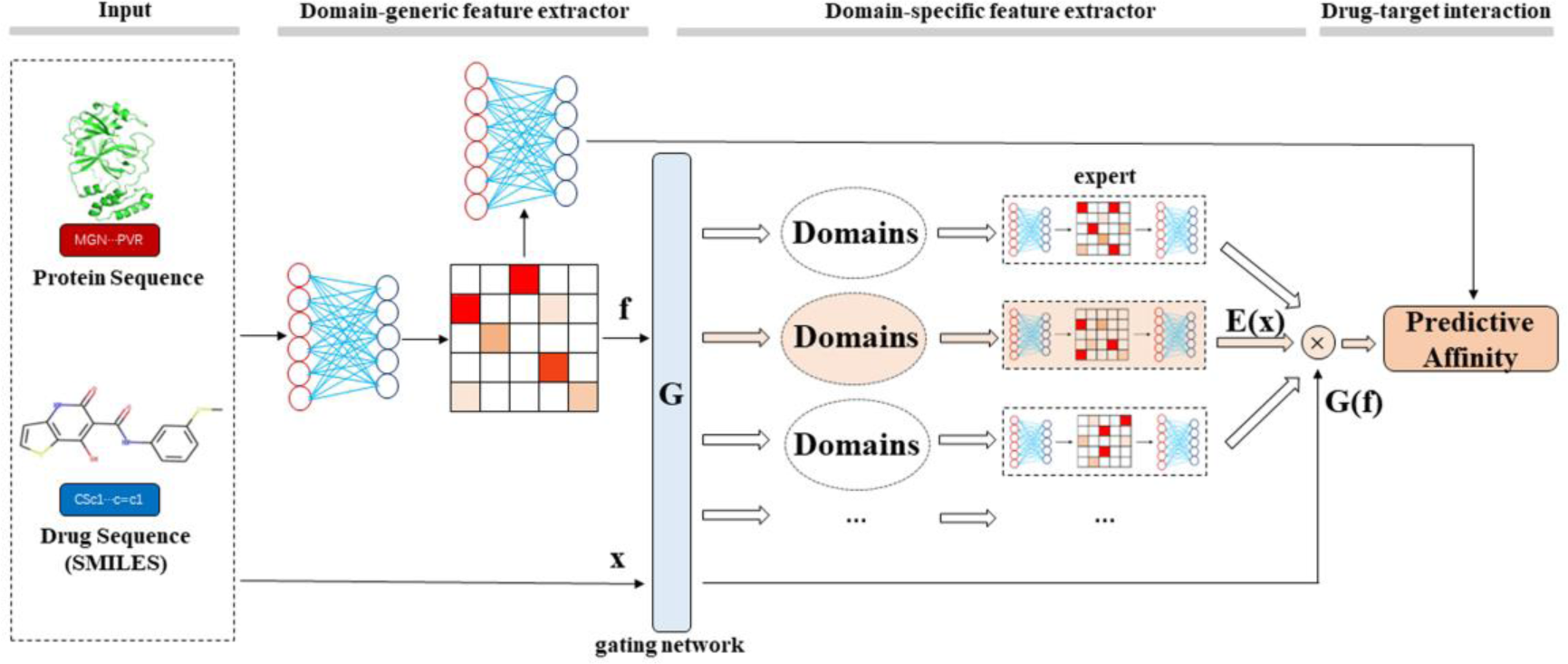
Overview of the E-DIS framework, which comprises ensembles of a domain-generic feature extractor and a domain-specific feature extractor. The gating network (G) routes the input data (X) to the corresponding experts based on the features learned by the domain-generic feature extractor.

### 2.3 domain-generic feature extractor

The domain-generic feature extractor in our framework can be any DTI prediction network that is trained using the entire training dataset. In our experiment, the domain-generic feature extractor encodes the input drug molecule and protein sequence separately. These encoded representation vectors are then concatenated to obtain the domain-generic feature *f*. The domain-generic feature serves multiple purposes in our framework. Firstly, it is utilized as input to various fully connected layers for predicting DTI. Additionally, it is also inputted into a gating network, which models the potential space of drug-target pairs as a combination of different domains. The domain-generic feature learned from the data is used as input to the gating network to determine which expert will be selected for a given input. For a given input x, the domain-generic feature *f* is

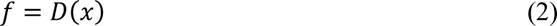

### 2.4 domain-specific feature extractor

MOE is trained on the different domains to capture domain-specific features. Unlike traditional deep learning models where the entire neural network participates in the computation for each input sample, MOE integrates multiple neural networks into a single task, with each network dedicated to a specific part of the dataset. Each of these neural networks is referred to as an expert. During both training and inference, an input dynamically activates a specific expert within the MOE. This activation is adaptively determined based on the input’s domain-generic feature and the expertise of the gating network. The final output of the MOE is a weighted combination of the outputs from each expert and the gating network. Consequently, each expert has a specific domain in which they excel and performs better than other experts.

The MOE framework in our experiments consists of a set of expert networks E1, · ·, En and a trainable gating network G. We have 3 experts in our experiments, but only need to evaluate one of them for each example. Each expert has specific learnable weights and is responsible for handling a different part of the dataset. While all experts are DTI prediction networks with identical architectures in our setting, it’s important to note that experts can have varying architectures as long as they accept inputs of the same size and produce outputs of the same size. This flexibility allows each expert to specialize in a particular domain.

The gating network plays a crucial role in routing the raw data to the appropriate experts, based on the features *f* learned by the domain-generic feature extractor. And the experts take the raw data as the input to avoid any potentially valuable features being discarded by the domain-generic feature extractor during the learning process. For a given raw data x, the output y of the MOE module is computed as follows:

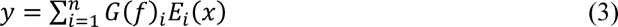

Where *f* is the feature learned by the domain-specific feature extractor, *G*(*f*)_i_ is the contribution of the i-th expert to the output vector, *E*_i_(*x*) is the output of the i-th expert network.

### 2.5 Metrics

For the regression task, we evaluated the models using mean square error (MSE, the smaller the better), concordance index (CI, the larger the better),[30] and r_m_ ^2^ index (the larger the better)[31] as performance metrics [32–34]. The MSE is a commonly used metric that measures the difference between the predicted and the actual values, with smaller values indicating higher prediction accuracy. The CI measures whether the predicted values maintain the same order as the real values. The MSE evaluates the prediction accuracy of models that output continuous values, while the CI evaluates the ranking performance. The r_m_ ^2^ index measures the external prediction performance of the model on a new dataset. In the classification task, we employed several performance metrics to evaluate the models. These included the area under the receiver operating characteristic curve (AUROC), the area under the precision-recall curve (AUPRC), accuracy, sensitivity, and specificity. AUROC and AUPRC measure the discrimination ability and precision of the models, respectively, with larger values indicating better performance. Accuracy represents the proportion of correctly classified samples, while sensitivity and specificity assess the model’s ability to correctly identify positive and negative samples, respectively.

## 3 Results and discussion

E-DIS were implemented in MindSpore [35]. We used the Adam optimizer [36] with the default momentum schedule. We employed two different experimental strategies: the in-domain setting and the cross-domain setting. In the in-domain setting, each experimental dataset was randomly divided into train set and test set according to a specific proportion. While the in-domain setting is a common experimental approach, it may not necessarily reflect the real-world performance of DTI prediction models in prospective predictions. Some studies have shown that results obtained through random splitting can be overly optimistic due to information leakage [6,10]. Furthermore, in real-world applications, the drug-target pairs that need to be predicted are usually unseen in the training data due to the vast chemical space. To imitate realistic scenario and further demonstrate the OOD generalization capability of E-DIS, we also adopted cross-domain setting. In the cross-domain setting, measures were taken to ensure that the structural information about drugs and targets did not leak into the test set. This provides a more realistic and challenging evaluation option for the models.

### 3.1 In-domain performance comparison regression tasks

In this study, we applied the proposed E-DIS to GraphDTA,[25] MGraphDTA[10] and DGraphDTA[34] and compared their performance with the benchmark models (DeepDTA,[32] WideDTA,[5] GraphDTA, DeepAffinity,[37] MGraphDTA and DGraphDTA). The results of DeepDTA, WideDTA, GraphDTA, DeepAffinity and MGraphDTA were taken from the reference[10]. We observed that the protein sequences in the majority of cases were characterized solely by amino acid types. However, DGraphDTA incorporated additional properties based on R groups such as polarity, weight, electrification, aromaticity, and others. To enhance the protein representation module of MGraphDTA, we incorporated these protein features while keeping the remaining aspects of the model consistent with MGraphDTA. To ensure a fair comparison, we used the same training and test sets as GraphDTA for the Davis and KIBA datasets. All experiments were repeated three times using different random seeds for robustness analysis.

Table 2 and Figure 2 showed the DTI prediction performance of E-DIS and the benchmark models on the Davis and KIBA datasets. The results indicated that incorporating additional protein representation information in MGraphDTA_P leads to a slight improvement in its DTI prediction performance compared to MGraphDTA. E-DIS_MGraphDTA_P_ has consistently outperformed other models in MSE and performed competitively in CI and r_m_ ^2^, despite did not using any information related to spatial structure. As can be seen from Figure 2, the prediction performance of all models significantly improved after applying the E-DIS method. The difference between the models before and after using the E-DIS method is statistically significant (95% confidence interval) according to the t-test (p < 0.05). In the KIBA dataset, GIN showed the most significant improvement before and after using the E-DIS method, with a 10.6% reduction in MSE. And MGraphDTA_P showed the most significant improvement in the Davis dataset, with a 6.7% reduction in MSE.

**Figure 2.**
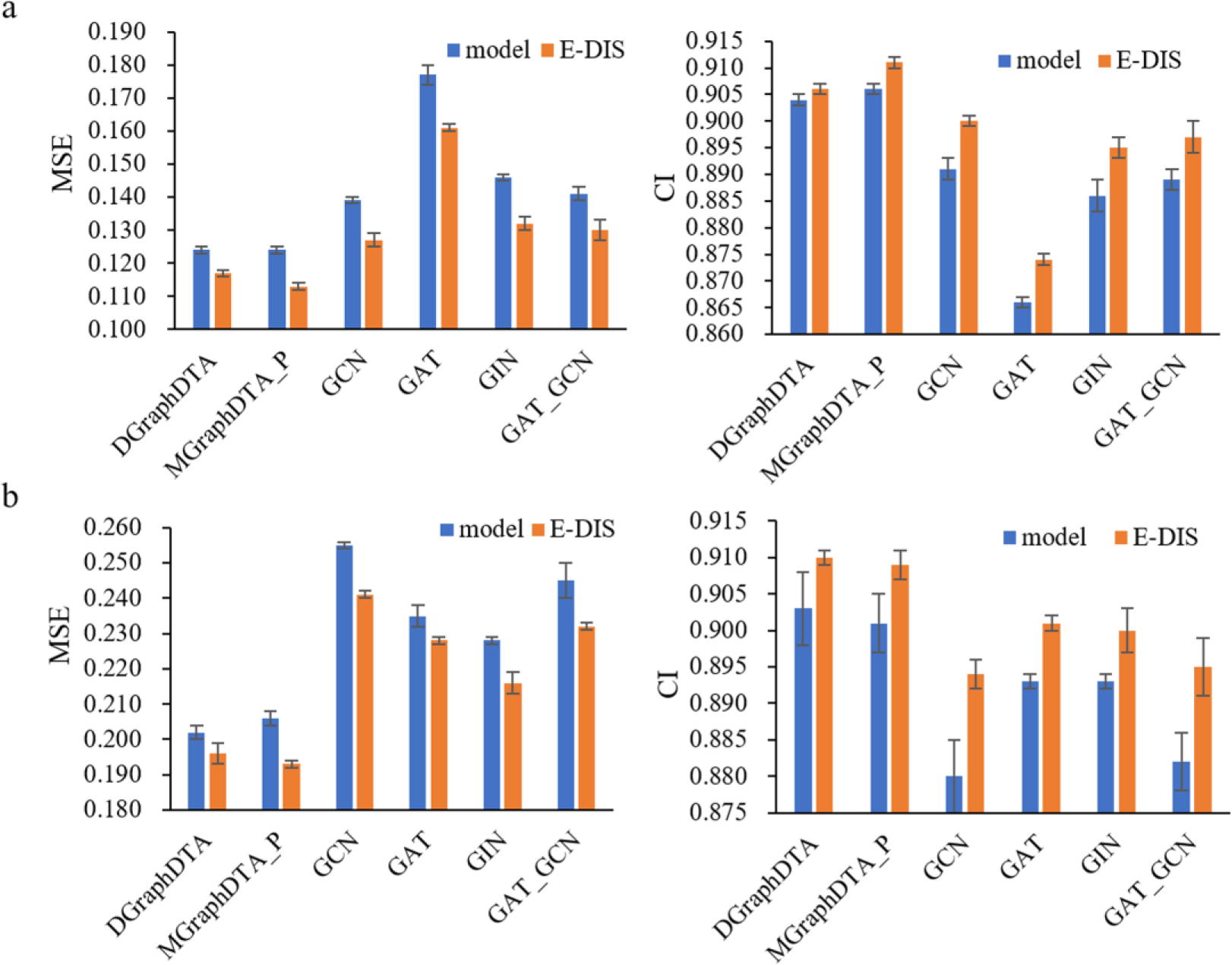
Performance comparison before and after using the E-DIS method on (a) KIBA and (b) Davis datasets with random split settings.

**Table 2.** Performance comparison on the Davis and KIBA datasets with random split settings (**Best**, <u>Second Best</u>).

| Datasets | Davis |  |  | KIBA |  |  |
| --- | --- | --- | --- | --- | --- | --- |
| Method | MSE | CI | $r_m^2$ index | MSE | CI | $r_m^2$ index |
| DeepDTA | 0.261 | 0.878 | 0.630 | 0.194 | 0.863 | 0.673 |
| WideDTA | 0.262 | 0.886 |  | 0.179 | 0.875 |  |
| GraphDTA | 0.229 | 0.893 |  | 0.139 | 0.891 |  |
| DeepAffinity | 0.253 | 0.900 |  | 0.188 | 0.842 |  |
| DGraphDTA | <u>0.202(0.002)</u> | <u>0.903(0.005)</u> | 0.708(0.007) | <u>0.124(0.001)</u> | 0.904(0.001) | 0.786(0.001) |
| MGraphDTA | 0.207(0.001) | 0.900(0.004) | <u>0.710(0.005)</u> | 0.128(0.001) | 0.902(0.001) | <u>0.801(0.001)</u> |
| MGraphDTA_P | 0.206(0.002) | 0.901(0.004) | 0.709(0.003) | <u>0.124(0.001)</u> | <u>0.906(0.001)</u> | 0.794(0.002) |
| E-DIS<sub>MGraphDTA_P</sub> | **0.193(0.001)** | **0.909(0.002)** | **0.732(0.003)** | **0.113(0.001)** | **0.911(0.001)** | **0.811(0.001)** |

#### classification tasks

The DTI prediction performance of E-DIS and the benchmark models was also compared for the classification task on the BindingDB and BIOSNAP datasets. We used the same dataset as DrugBAN under the random split setting. The results of GraphDTA, MolTrans and DrugBAN were taken from the references[6], and the rest of the experiments were repeated three times to calculate the mean and standard deviation of the prediction results.

Table 3 and Figure 3 showed the DTI prediction performance of E-DIS and the benchmark models on the BindingDB and BIOSNAP datasets. It can be observed that the DTI prediction performance of all models improved after applying the E-DIS method, indicating that utilizing both domain-generic features and domain-specific features contributed to enhancing the model’s prediction performance. E-DIS_DrugBAN_ consistently outperformed other models in two datasets. The AUROC for E-DIS_DrugBAN_ on BindingDB and BIOSNAP were 0.971(0.000), 0.921(0.001), respectively. At the same time, it is worth noting that models that do not use E-DIS method achieved similar and excellent performance on both datasets, and there was no significant difference between each model. This may be attributed to the overoptimistic performance estimation under the random split setting caused by information leakage, which does not accurately reflect the real-world performance of the models in prospective prediction scenarios.

**Figure 3.**
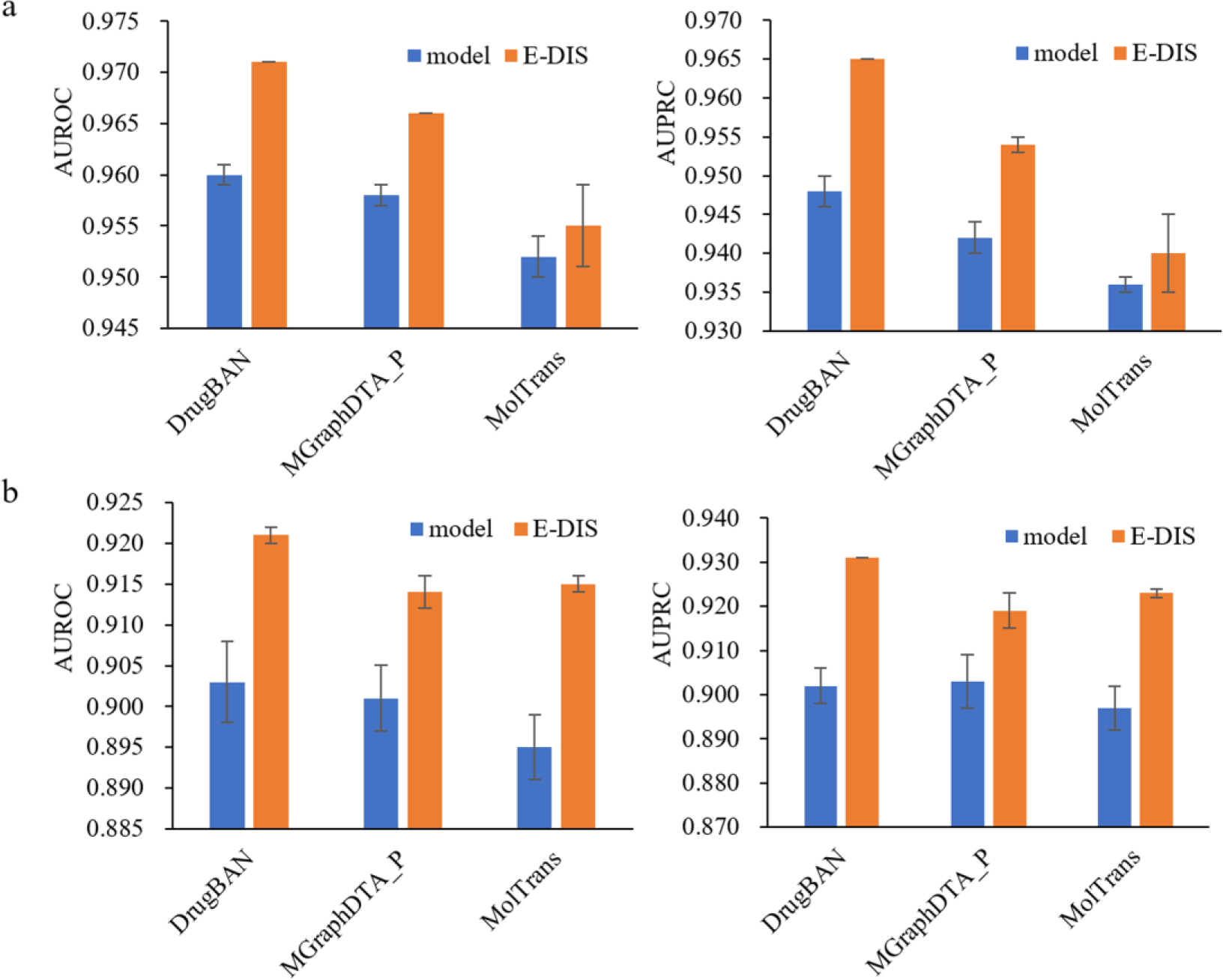
Performance comparison before and after using the E-DIS method on (a) BindingDB and (b) BIOSNAP datasets with random split settings.

**Table 3.**
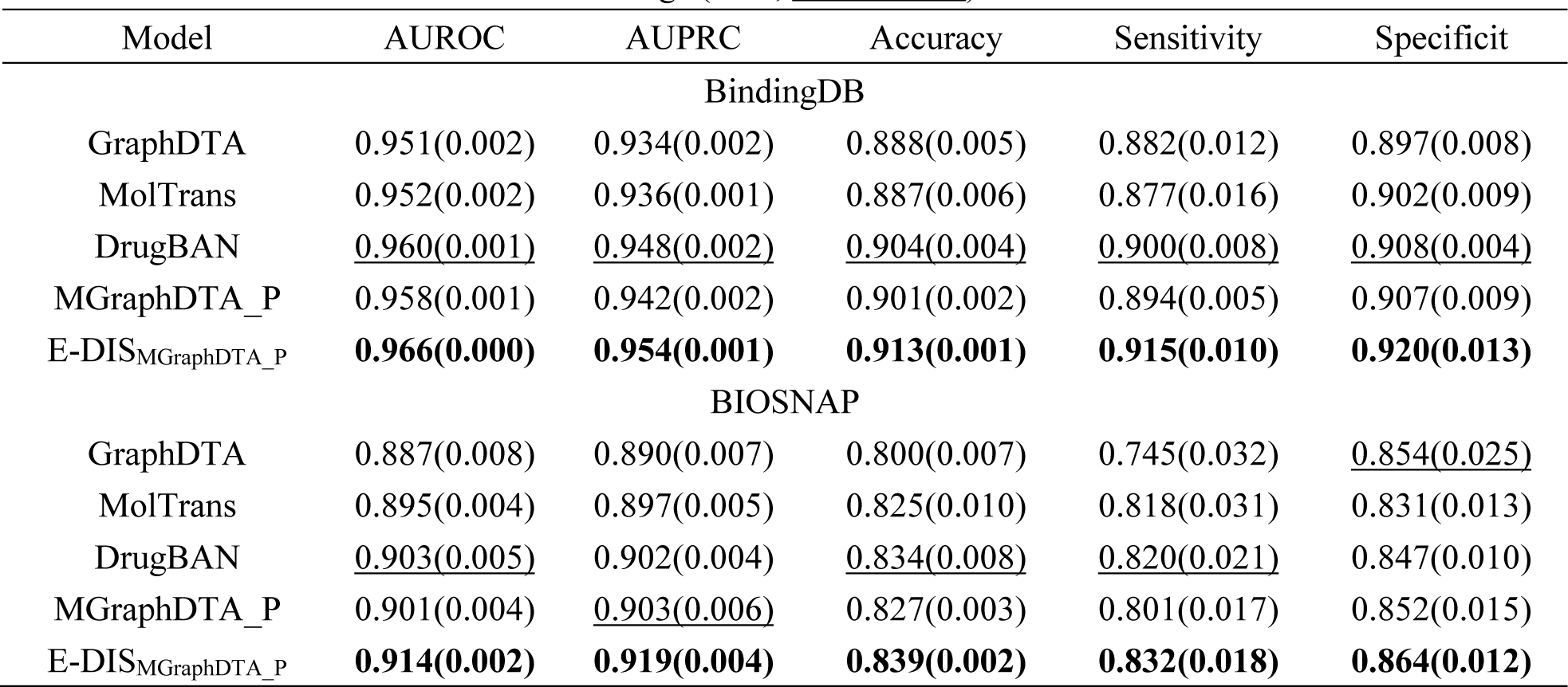
Performance comparison on the BindingDB and BIOSNAP datasets with random split settings (**Best**, <u>Second Best</u>).

### 3.2 Cross-domain performance comparison unseen drug/target split

Additional experiments were conducted using different splitting strategies on the BindingDB and BIOSNAP datasets to ensure that the structural information about drugs and targets does not leak into the test set. In this approach, 10% of the drugs/proteins were randomly selected as the test set, and all associated drugs/proteins were removed from the training set. This ensured that the test set contained unseen drugs or proteins that were not present in the training set. The performance of the baseline models (MGraphDTA_P, MolTrans, and DrugBAN) was compared before and after applying the E-DIS method. For a fair comparison, all models shared the same training and test sets, and all experiments were repeated three times.

Figure 4 showed the results of three experiments conducted under the unseen drug/target split settings. A significant degrade in performance can be observed compared to the results obtained under the random split setting shown in Table 4. None of the models consistently maintained optimal performance across all split settings, indicating the greater difficulty of the split settings that involve unseen drugs/targets. Under the unseen drug/target split setting, the prediction performance of the three models showed improvement to varying degrees after applying the E-DIS method. Particularly, MGraphDTA_P and MolTrans exhibited highly significant differences (p < 0.01) in performance before and after using the E-DIS method under the unseen drug split, while DrugBAN also showed significant differences (p < 0.05). E-DIS_MGraphDTA_P_ achieved a 7.9% increase in AUROC compared to MGraphDTA_P in the BindingDB dataset and a 5.1% increase in the BIOSNAP dataset. In general, the majority of models (17 out of 18) demonstrated significantly improved performance after using E-DIS in the two datasets across the three split methods. These results highlight that E-DIS can enhance the prediction accuracy of DTI prediction models for unseen drugs/targets in the training set.

**Figure 4.**
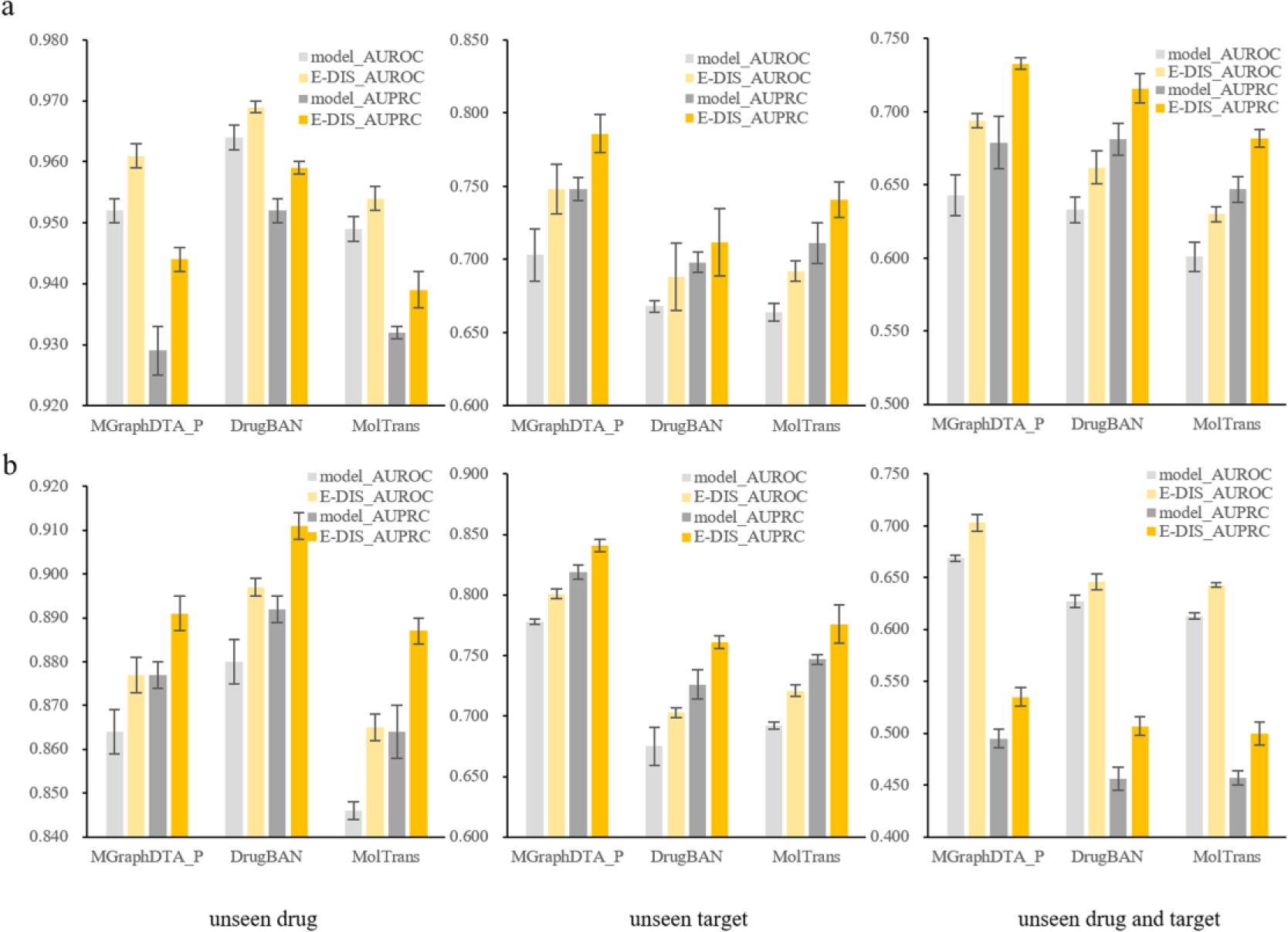
Performance comparison before and after using the E-DIS method on the (a) BindingDB and (b) BIOSNAP datasets with unseen drug, unseen target, unseen drug and target split settings.

**Table 4.** Performance comparison before and after using the E-DIS method on the BindingDB and BIOSNAP datasets with unseen drug/target split settings. (**Best**, <u>Second Best</u>).

| Method | BindingDB |  | BIOSNAP |  |
| --- | --- | --- | --- | --- |
|  | AUROC | AUPRC | AUROC | AUPRC |
| unseen drug |  |  |  |  |
| MGraphDTA_P | 0.952(0.002) | 0.929(0.004) | 0.864(0.005) | 0.877(0.003) |
| E-DIS<sub>MGraphDTA_P</sub> | 0.961(0.002) | 0.944(0.002) | 0.877(0.004) | 0.891(0.004) |
| MolTrans | 0.949(0.002) | 0.932(0.001) | 0.846(0.002) | 0.864(0.006) |
| E-DIS<sub>MolTrans</sub> | 0.954(0.002) | 0.939(0.003) | 0.865(0.003) | 0.887(0.003) |
| DrugBAN | <u>0.964(0.002)</u> | <u>0.952(0.002)</u> | <u>0.880(0.005)</u> | <u>0.892(0.003)</u> |
| E-DIS<sub>DrugBAN</sub> | **0.969(0.001)** | **0.959(0.001)** | **0.897(0.002)** | **0.911(0.003)** |
| unseen target |  |  |  |  |
| MGraphDTA_P | <u>0.703(0.018)</u> | <u>0.748(0.008)</u> | <u>0.778(0.002)</u> | <u>0.819(0.006)</u> |
| E-DIS<sub>MGraphDTA_P</sub> | **0.748(0.017)** | **0.786(0.013)** | **0.801(0.004)** | **0.841(0.005)** |
| MolTrans | 0.664(0.006) | 0.711(0.014) | 0.692(0.003) | 0.747(0.004) |
| E-DIS<sub>MolTrans</sub> | 0.692(0.007) | 0.741(0.012) | 0.721(0.005) | 0.776(0.016) |
| DrugBAN | 0.668(0.004) | 0.698(0.007) | 0.675(0.016) | 0.726(0.012) |
| E-DIS<sub>DrugBAN</sub> | 0.688(0.023) | 0.712(0.023) | 0.703(0.004) | 0.761(0.005) |
| unseen drug and target |  |  |  |  |
| MGraphDTA_P | 0.643(0.014) | 0.679(0.018) | <u>0.669(0.003)</u> | 0.495(0.009) |
| E-DIS<sub>MGraphDTA_P</sub> | **0.694(0.005)** | **0.733(0.004)** | **0.703(0.008)** | **0.535(0.009)** |
| MolTrans | 0.601(0.010) | 0.647(0.009) | 0.613(0.003) | 0.457(0.007) |
| E-DIS<sub>MolTrans</sub> | 0.630(0.005) | 0.682(0.006) | 0.643(0.002) | 0.500(0.011) |
| DrugBAN | 0.633(0.009) | 0.681(0.011) | 0.627(0.006) | 0.456(0.011) |
| E-DIS<sub>DrugBAN</sub> | <u>0.662(0.011)</u> | <u>0.716(0.010)</u> | 0.646(0.008) | <u>0.507(0.009)</u> |

**Table 5.** Performance comparison of E-DIS, BAT and CDAN on the BindingDB and BIOSNAP datasets with cluster-based split (**Best**, <u>Second Best</u>).

| Method | BindingDB |  | BIOSNAP |  |
| --- | --- | --- | --- | --- |
|  | AUROC | AUPRC | AUROC | AUPRC |
| MolTrans | 0.565(0.004) | 0.520(0.011) | 0.614(0.008) | 0.607(0.010) |
| BAT<sub>MolTrans</sub> | 0.594(0.007) | <u>0.605(0.009)</u> | 0.628(0.003) | 0.649(0.037) |
| CDAN<sub>MolTrans</sub> | 0.606(0.012) | 0.566(0.027) | 0.625(0.004) | 0.638(0.020) |
| E-DIS<sub>MolTrans</sub> | 0.618(0.004) | 0.601(0.042) | 0.656(0.011) | 0.691(0.017) |
| DrugBAN | 0.571(0.013) | 0.526(0.007) | 0.624(0.009) | 0.605(0.009) |
| BAT<sub>DrugBAN</sub> | 0.587(0.015) | 0.573(0.021) | 0.648(0.008) | 0.670(0.011) |
| CDAN<sub>DrugBAN</sub> | 0.604(0.014) | 0.558(0.020) | 0.682(0.012) | 0.721(0.011) |
| E-DIS<sub>DrugBAN</sub> | 0.611(0.008) | **0.609(0.027)** | 0.660(0.006) | 0.690(0.010) |
| MGraphDTA_P | 0.601(0.009) | 0.549(0.012) | 0.690(0.023) | 0.683(0.021) |
| BAT<sub>MGraphDTA_P</sub> | <u>0.624(0.021)</u> | 0.568(0.021) | <u>0.714(0.016)</u> | <u>0.724(0.015)</u> |
| CDAN<sub>MGraphDTA_P</sub> | 0.617(0.014) | 0.585(0.025) | 0.693(0.003) | 0.696(0.004) |
| E-DIS<sub>MGraphDTA_P</sub> | **0.630(0.004)** | 0.601(0.011) | **0.721(0.004)** | **0.732(0.004)** |

#### cluster-based split

Furthermore, we also adopted a more stringent cluster-based splitting strategy to evaluate the predictive performance of the model on data with unknown distributions. The same source and target domains as used in DrugBAN were selected under the cluster-based split setting. The drugs and targets were clustered using ECFP4 fingerprint and pseudo amino acid composition, respectively. Drug-target pairs between 60% of the drug clusters and 60% of the target clusters were considered as the source domain data, while the remaining clusters were considered as the target domain data. This ensured that the compounds in the source and target domains had different structures, as the single-linkage clustering method was used to maintain a sufficient distance between any two clusters.

Table5 and Figure 5 showed the performance of E-DIS compared to BAT and CDAN, which were advanced methods for solving out-of-distribution generalization problem under the cluster-based split setting. It was observed that models using the E-DIS method (E-DIS_MODEL_) significantly outperformed the models without using E-DIS (MODEL) in both the BindingDB and BIOSNAP datasets (p < 0.01). The average AUROC of E-DIS_MODEL_ increased by 7.1% and 5.7% compared to MODEL in the BindingDB and BIOSNAP datasets, respectively. Among all models, E-DIS_MGraphDTA_P_ consistently outperformed the others in AUROC and performed competitively in AUPRC. Furthermore, the performance of most models using the E-DIS method was better than those using BAT or CDAN. Notably, E-DIS did not use any information about the target domain during training, while both BAT and CDAN relied on information from unlabeled data in the target domain. These results highlight that the E-DIS method achieved comparable or even superior performance compared to BAT and CDAN in solving the out-of-distribution generalization problem, showcasing the benefits of domain-generic features and domain-specific features for model generalization.

**Figure 5.**
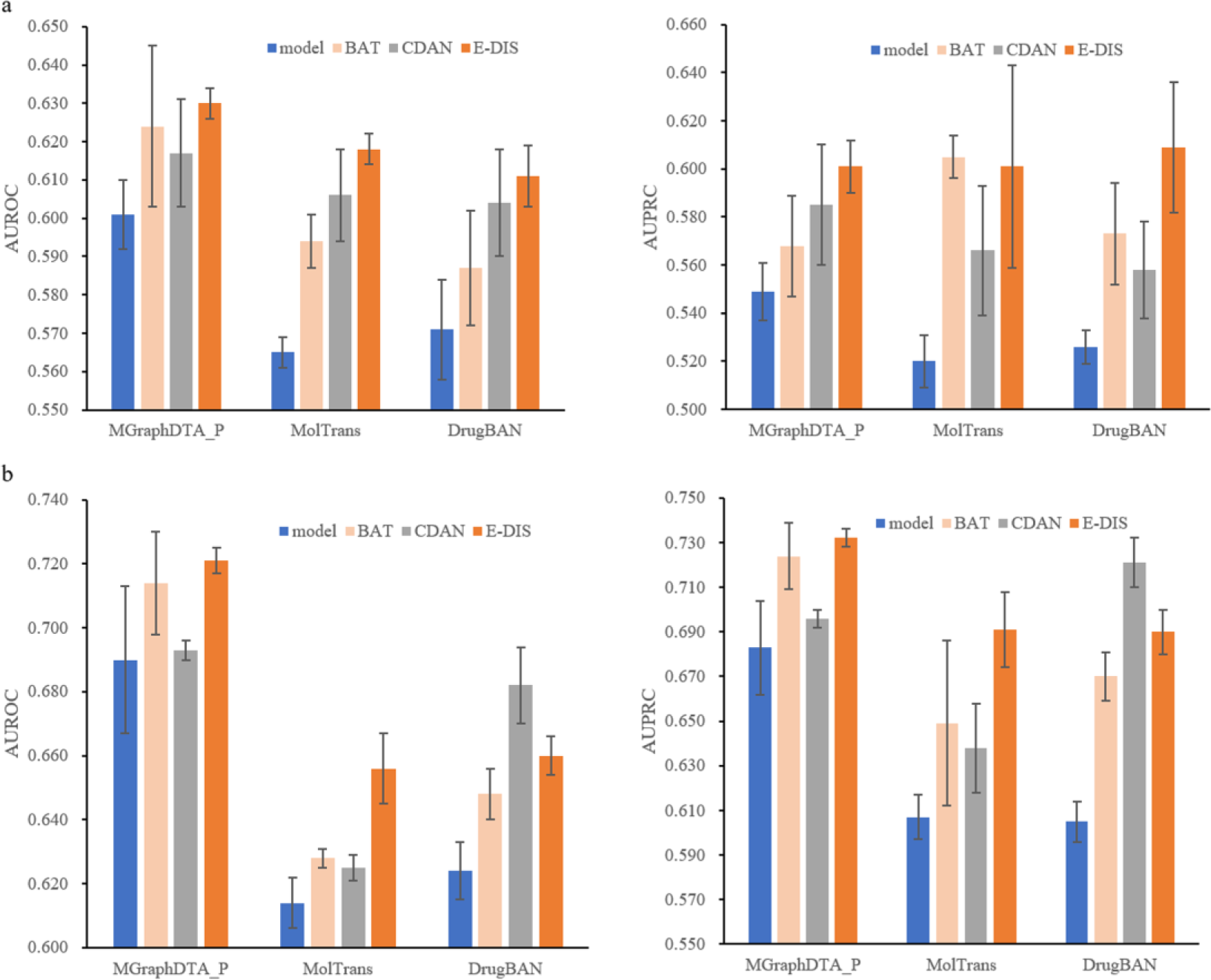
Performance comparison of E-DIS, BAT and CDAN on the (a) BindingDB and (b) BIOSNAP datasets with cluster-based split settings.

### 3.3 Interpretation and visualization

In order to explore whether the two extractors in E-DIS learn different features, we visualized the importance of atoms in the molecules, as shown in Figure 6. We calculated each atom’s contribution to the final predicted result using the Grad-AAM method,[10] a visual interpretation technique that use the gradient information from the last graph neural network layer. And we then generated a probability map using RDkit. The visualization results demonstrated that both extractors were able to detect essential features related to hydrogen bond, hydrophilic, hydrophobic, π-π interactions and other important factors involved in drug-target binding. Furthermore, it was observed that the contributions of atoms in the molecules differed between the two extractors, indicating that they focused on different types of interactions. Specifically, the domain-generic feature extractor primarily emphasized hydrophilic, hydrophobic and π-π interactions, while the domain-specific feature extractor placed more emphasis on hydrogen bond interactions.

**Figure 6.**
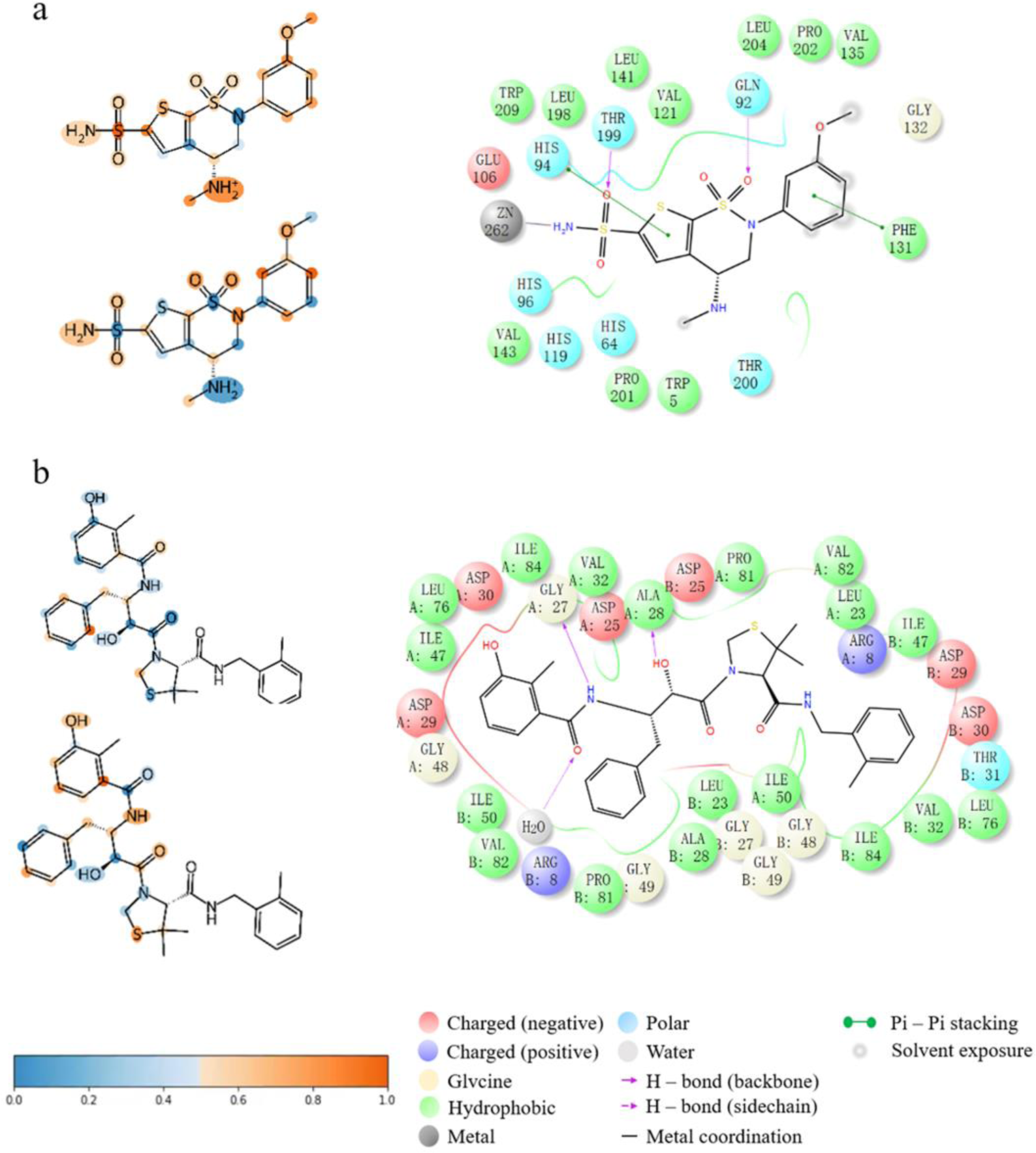
Importance visualization of co-crystalized ligands (a: 1BNM, b: 1KZK) from Protein Data Bank (PDB). The upper left side of each panel shows the importance visualization of atoms in domain-generic feature extractor. The bottom left side of each panel shows the importance visualization of atoms in domain-specific feature extractor. The right side of each panel shows the 2D interaction visualization extracted from the complex. At the bottom right, the legend panel for the drug-target interaction maps is displayed.

These findings suggest that the two extractors in E-DIS learn complementary features, with each extractor specializing in different aspects of drug-target interactions. This supports the notion that the combination of domain-generic and domain-specific features in E-DIS enables a comprehensive understanding of the complex interactions between drugs and targets.

## 4 Conclusions

The accurate prediction of DTI is a crucial step for the field of new drug design and development. Although deep learning models have shown promise in DTI prediction, their limited generalization ability hinders their practical application. In this paper, we propose a versatile ensemble of models called E-DIS, which aims to capture both domain-generic features and domain-specific features. By incorporating diverse domain features, E-DIS mitigates the risk of relying solely on one feature and thus improves its generalizability to novel drug-target pairs. The experimental results demonstrate that E-DIS consistently outperforms other models in DTI prediction tasks, both in in-domain and cross-domain settings. Furthermore, E-DIS exhibits competitive performance in domain generalization, comparable to related approaches. In our experiment, the experts in different domains are homogeneous networks with identical architectural units that differ only in parameters. However, in future research, experts with varying architectural units could be explored to better accommodate the diversity of inputs. Importantly, E-DIS is algorithmically flexible and can be easily extended to other fields beyond DTI prediction. This versatility makes it a promising approach for addressing generalization challenges in various domains.

## Key Points

1. E-DIS is a versatile framework for predicting drug-target interactions that enables training different extractors to capture domain-generic and domain-specific features respectively.
2. E-DIS enhances the generalizability to novel drug-target pairs by training multiple experts to capture and align domain-specific information from the training domains without accessing any data from unseen domains.
3. To demonstrate the efficacy of our proposed E-DIS framework, we extensively evaluate it on four benchmark datasets under both in-domain and cross-domain settings. The majority of models showed significantly improved performance after using E-DIS (p < 0.05) and perform competitively with advanced approaches in domain generalization.

## Data and code availability

All data used in this paper are publicly available and can be accessed at https://github.com/hkmztrk/DeepDTA/tree/master/ data for the Davis and KIBA datasets, https://github.com/peizhenbai/DrugBAN/tree/main/datasets data for the BindingDB and BIOSNAP datasets. All codes of E-DIS are available at https://www.mindspore.cn/ecosystem/libraries.

## Acknowledgments

This work was supported by the National Key R&D Program of China under Grant Nos. 2022ZD0117804 and 2021ZD0110400. We thank the Supercomputing Center of Lanzhou University for providing high-performance computing resources.

## Author Contributions

S.L., J.Y., Z.W., N.N., M.C., J.Z., X.Y. and H.L. conceived the project. S.L., J.Y., Y.L., C.X., J.Z., X.Y. and H.L. designed and implemented the experiments. S.L., J.Y., M.C., Y.L., C.X., Y.D., J.Z., X.Y. and H.L. analyzed the results. All authors co-wrote and reviewed the manuscript.

